# The third symbiotic partner of the volcano lichen *Cladonia vulcani* Savicz drove adaptation to an extreme environment

**DOI:** 10.1101/2023.04.05.535799

**Authors:** Mieko Kono, Yohey Terai

**Affiliations:** Research Center for Integrative Evolutionary Science, SOKENDAI (The Graduate University for Advanced Studies), Shonan Village, Hayama, Kanagawa, 240-0193, Japan

## Abstract

Chemosynthetic symbioses between sulfur-oxidizing bacteria and aquatic eukaryotes have been discovered globally in sulfide-rich environments, notably deep-sea hydrothermal vents, cold seeps, and sulfidic cave systems. However, to the best of our knowledge, such chemosymbiotic lifestyles have not been reported from terrestrial eukaryotes. Here we report that the volcano lichen *Cladonia vulcani* Savicz ubiquitously associates with a single bacterial species that could potentially use hydrogen sulfide as a source of energy. We identified sequences of the bacterium in all 27 samples collected from five geothermal areas across Japan with cellular abundance comparable to the fungal partner. The assembled bacterial genome contained genes involved in sulfur oxidation. The stable association with a potential sulfur-oxidizer is likely to represent an obligate tripartite symbiotic system consisting of fungal, algal, and bacterial partners that has enabled adaptation to the extreme environment.

## Introduction

Hydrogen sulfide is famous for its toxic properties primarily inhibiting cytochrome *c* oxidase in the mitochondrial respiratory chain (1). However, in deep-sea hydrothermal vent environments, chemoautotrophic sulfur-oxidizing bacteria use hydrogen sulfide or other reduced sulfur compounds in hydrothermal vent fluids to fuel carbon fixation that sustains the entire vent ecosystem (2-6). Complex megafauna found in one of the most extreme environments on Earth were formed through symbiotic interactions between sulfur-oxidizing bacteria and macroinvertebrates that facilitated adaptation of host-symbiont systems (7-14). Such chemosynthetic lifestyles were later reported from other sulfide-rich environments such as cold seeps (15, 16) and sulfidic cave systems (17, 18), but none from terrestrial geothermal areas albeit the abundant emission of hydrogen sulfide gas.

Geothermal areas with volcanic steam vents that constantly emit significant hydrogen sulfide gas typically present desolated open fields with very limited vegetation. The volcano lichen *Cladonia vulcani* Savicz is one of the few organisms that can thrive in such areas commonly forms extensive covers on ground beside active vents (19). Recently the classical view of lichens as two partners symbiosis, composed of one fungus (mycobiont) and one alga/cyanobacterium (in rare cases alga and cyanobacterium) (photobiont), has been challenged by studies that reported ubiquity and structured composition of lichen-associated bacterial communities (20-33). Bacteria are presumed to have supportive roles that can enhance fitness of lichens under extreme environments (30, 32). Therefore, it is not difficult to imagine that *C. vulcani* has become adaptive to sulfide-rich environments by recruiting a third microbial symbiotic partner.

The objective of this study was to identify universal associates of *C. vulcani* besides the fungal and algal partners that may have contributed to the adaptation to sulfide-rich environments. For this purpose, we *de novo* assembled the genomes of the fungal and algal partners of *C*.*vulcani* and obtained metagenomic data from 27 samples collected from five volcanic areas known for active hydrogen sulfide gas emissions in Japan. The metagenomic datasets were searched for sequences universally existing in the samples and functions of a third symbiotic partner were predicted from assembled genomic sequences.

## Methods

### Sample collection and hydrogen sulfide measurement

*Cladonia vulcani* was sampled from five geothermal areas in three islands of Japan (HOKKAIDO. Pref. Hokkaido: Noboribetsu spa. HONSHU. Pref. Aomori: Mt. Hakkoda, Pref. Gunma: Kusatsu spa., Pref. Kanagawa: Owakudani. KYUSHU: Pref. Kumamoto: Mt. Aso) with permissions from National Parks administering the areas. At each locality, podetia were picked using sterilized tweezers and immediately put into RNAlater (Thermo Fisher Scientific Inc.). The samples were stored at -20 °C until use. In each geothermal area, we measured the concentration of hydrogen sulfide in ambient air using 4LB (GASTECH CORPORATION) hydrogen sulfide detector tubes, following the manufacturer’s instructions. The concentration of hydrogen sulfide was monitored during the sampling in Hakkoda using a gas detector (XS-2200, New Cosmos Electric Co., LTD.).

### Short-read genome sequencing on the Illumina platform

Metagenomic DNA extraction was performed using a single podetium per extraction (occasionally two podetia were used to ensure the amount of DNA required in library preparation) with at least three biological replicates from each sampling locality. DNA of the fungal and algal symbionts were extracted from independent cultures isolated from lichen samples collected in Noboribetsu. The lichen and fungal samples were homogenized with stainless steel beads (SUB-30, TOMY), whereas algal samples were frozen in liquid nitrogen and homogenized using Automill (TK-AM7-24, Tokken Inc.). DNA was extracted using DNeasy Plant Mini kit (QIAGEN) following the manufacturer’s instructions. DNA libraries were constructed using NEBNext Ultra II DNA Library Prep Kit for Illumina and NEBNext Multiplex Oligos for Illumina (New England Bio Labs) following the manufacturer’s instructions. The libraries were sequenced on the Illumina Hiseq X platform with 150 bp paired-end reads (125 bp paired-end reads for the samples from Noboribetsu).

### Long-read genome sequencing on the Oxford Nanopore MinION platform

High-molecular weight DNA was extracted from podetia of Noboribetsu samples using buffers in DNeasy Blood & Tissue Kit and Genomic-tip 20/G (QIAGEN) with lysis time extended to 5 hours. A long-read DNA library was constructed using Rapid Sequencing Kit (SQK-RAD004, Oxford Nanopore Technologies) following the manufacturer’s protocol. The library was loaded on a R9.4.1 flow cell primed using Flow Cell Priming Kit (EXP-FLP002) and sequenced on a MinION device following the manufacturer’s instructions. MinKNOW was used to drive the sequencing including the data acquisition.

### Assembly of fungal and algal genomes

The reads were subjected to genome assemblies after removal of adaptor sequences and low-quality reads. The genome of the mycobiont was assembled by using CLC Genomic Workbench (QIAGEN) in a combination of Illumina and Nanopore reads. First, the Illumina short-reads derived from the fungal independent culture were mapped on the Nanopore long-reads to extract consensus sequences. Then, the short-reads and the consensus sequences were *de novo* assembled with a word size of 64. The genome of the photobiont was *de novo* assembled using Illumina short-reads from the independent algal culture with default settings of CLC Genomic Workbench. Completeness of the genomes was assessed using Benchmarking Universal Single-Copy Orthologs (BUSCO) (34).

### Identification of unknown symbiotic partners

The short-reads of three (out of 15) biological replicates from Noboribetsu (S1 reads) were *de novo* assembled into draft metagenomic contigs using CLC Genomic Workbench with default settings. Then, the short-reads of 27 lichen samples were mapped to the draft contigs and the number of reads mapped to a contig was plotted against the contig length (Figure 2). BlastN searches against the NCBI nr/nt database were performed on contigs aligned in two steep slopes in the plots.

**Figure 1.**
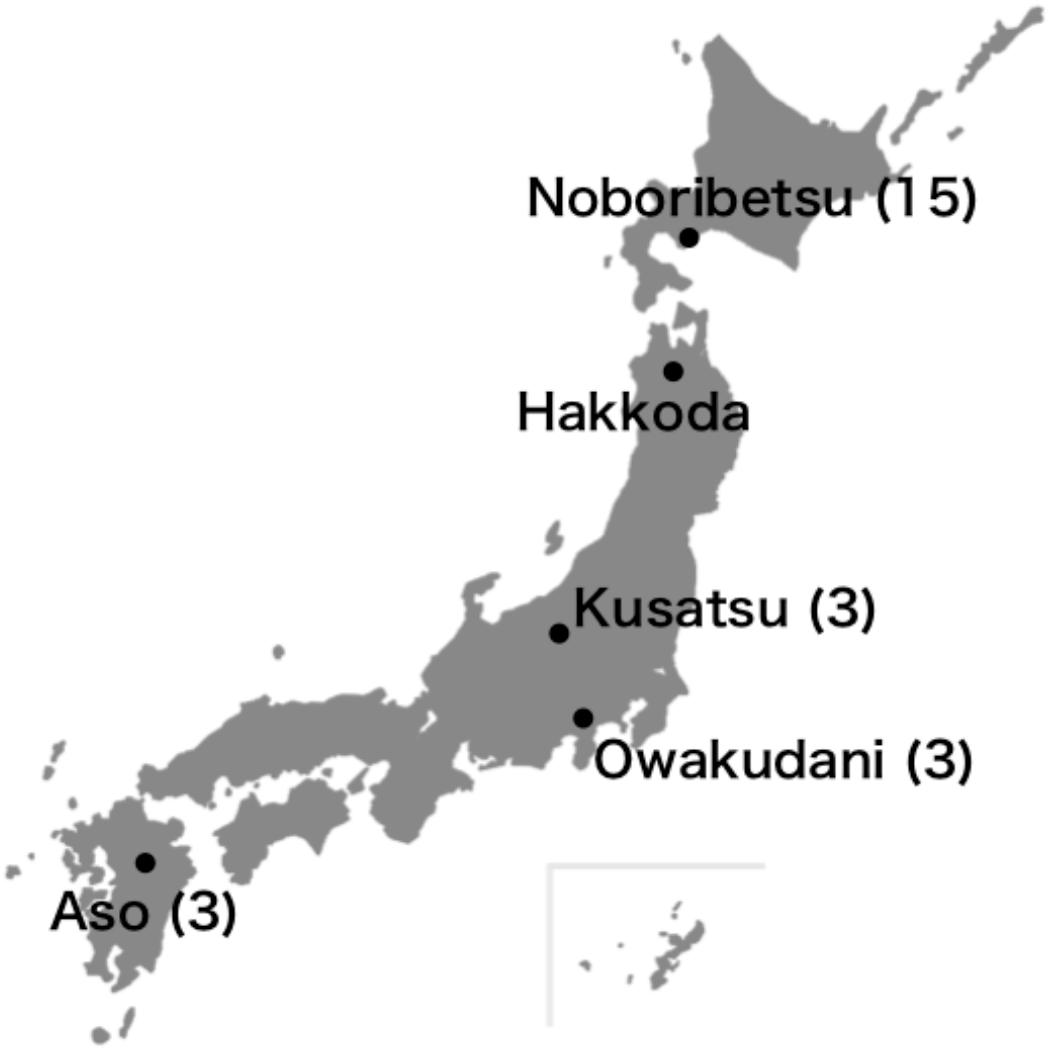
The samples used in this study were collected from the three islands of Japan. The number of samples collected in each locality is shown in the brackets.

**Figure 2.**
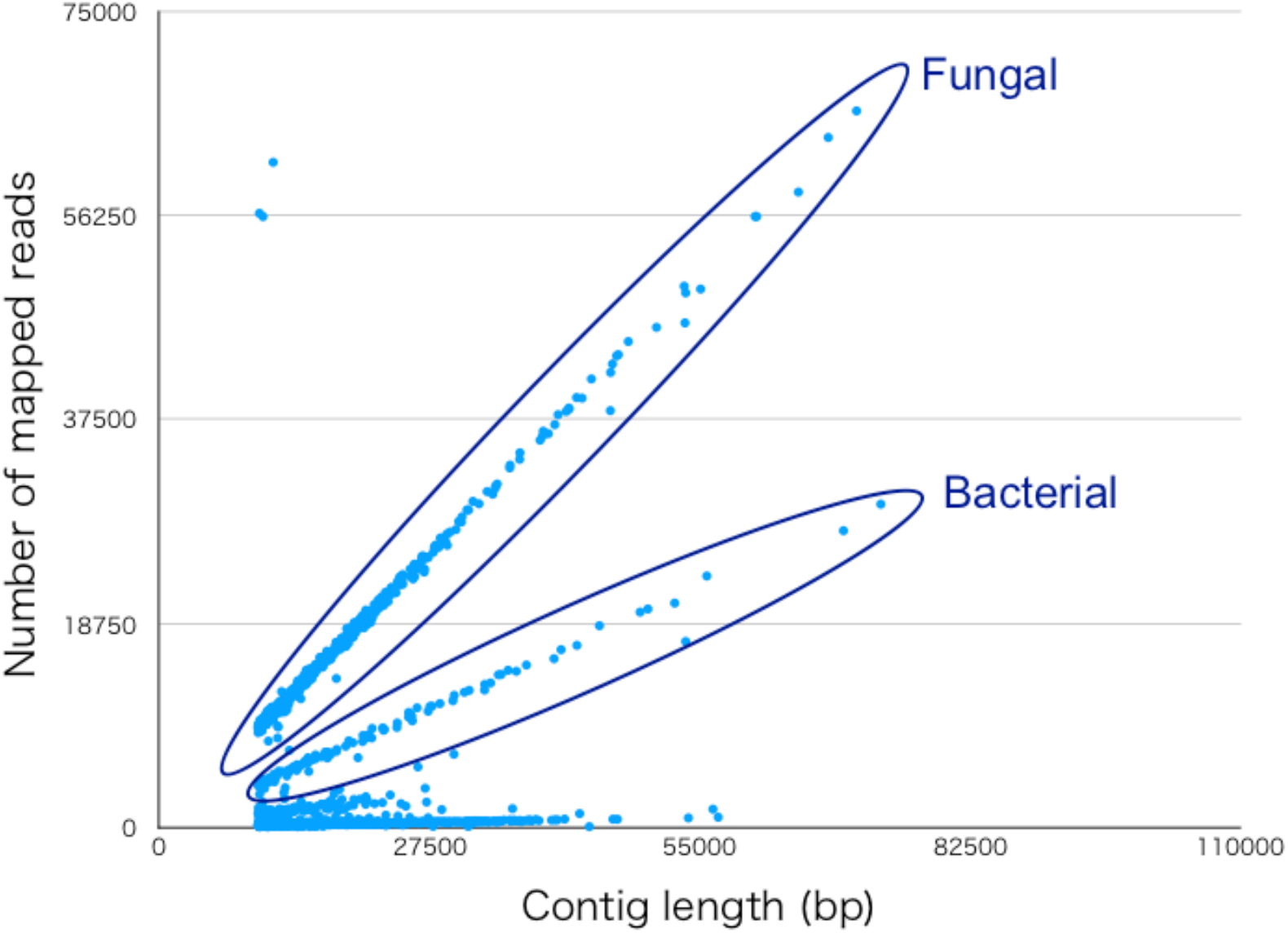
The number of metagenomic reads mapped to the draft metagenomic contigs (y-axis) is plotted against the contig length (x-axis). The plot of a replicate from Noboribetsu (contigs ≥ 10 kb) is shown in this figure. Two groups of contigs show higher-coverage than other contigs with similar lengths. BLASTN searches indicated that the group with larger slope is similar to fungal and smaller slope is similar to bacterial sequences.

### Genome assembly of the third bacterial partner (LFB_Cvu01)

The draft metagenomic contigs (≥ 10 kb) were subjected to DIAMOND blastX searches (v2.0.4.142) with a sensitive mode. The contigs were binned according to taxonomic groups by using MEGAN. Metagenomic contigs binned as *Proteobacteria* were used as template contigs to map the following sequences and extract consensus sequences: (i) Nanopore long-reads polished with the S1 reads (threshold = 2), (ii) Consensus sequences of the S1 reads mapped to the contigs assembled from S1 reads that were un-mapped to the draft metagenomic contigs binned as *Ascomycota* and *Viridiplantae* (threshold = 10). Consensus sequences of the long-reads mapped to the CDS of *Acidiphilium multivorum* AIU301 (downloaded from NCBI, GCA_000202835.1) were also extracted (threshold = 2) (iii). The sequences (i, ii, iii) were *de novo* assembled as a draft bacterial genome. Then, the short-reads from the five localities were combined and mapped to the draft bacterial genome to distinguish contigs of the third symbiotic partner by the normalized number of mapped reads (number of mapped reads/contig length). The S1 reads and long-reads were mapped to the contigs with high normalized numbers of mapped reads (≥ 10) and consensus sequences were extracted (threshold = 10 and 2, respectively) (iv). Finally, the sequences (iii) and (iv) were *de novo* assembled as genomic sequences of the third bacterial partner (LFB_Cvu01).

### Relative amount of LFB_Cvu01 in *C. vulcani*

Cellular abundance of LFB_Cvu01 in a lichen thallus was predicted from the ratio of genome coverage of LFB_Cvu01 to the mycobiont, calculated as follows, [(The number of mapped reads) x (the average length of mapped reads)] / (The total length of the genome).

### Phylogenetic analysis

Phylogenetic trees of the mycobiont and LFB_Cvu01 were constructed from the 27 metagenomic datasets. The short-reads were trimmed and mapped to the genomes using CLC Genomic Workbench. Duplicated read alignments were marked using MarkDuplicates algorithm implemented in GATK v4.2 (https://gatk.broadinstitute.org/hc/en-us. Variants [single nucleotide polymorphisms (SNPs) and insertions and deletions] were called using the HaplotypeCaller algorithm in GATK v4.2 in “-ERC GVCF” mode to produce GVCF files (35). The GVCF files were combined into a single file and passed to GATK GenotypeGVCFs for joint genotyping. Genotypes were filtered using VCFtools (36) with parameters “-remove-indels --max-missing 1 --minGQ 8 --minDP 5 --maxDP 1000”. The genotype table was converted to a Phylip (Interleaved) file using Tassel5 (https://tassel.bitbucket.io) after filtering with “Minimum Count 27, Max Heterozygous Proportion 0.8”. The first 10 kb of the aligned sequences were used to find the best substitution model for phylogenetic analysis using MEGA ver. X (37). Phylogenetic trees were constructed using PhyML with the best model (GTR) and a bootstrap analysis (100 replicates) (38).

### Identification of the genes involved in sulfur metabolism

Genes involved in sulfur metabolism were searched for the LFB_Cvu01 genome using tBLASTX (39) and DFAST (40). Query sequences for the tBLASTX searches were downloaded from the sulfur metabolism pathway in the KEGG database (41).

## Results

### Dependency on hydrogen sulfide gas

We collected thalli of *Cladonia vulcani* Savicz near solfataras (volcanic steam vents constantly emitting sulfurous gases) in five localities across Japan (Figure 1). In our surveys, the hydrogen sulfide gas was detected in all localities showing the concentration of over a hundred ppm by solfataras. Populations of *C. vulcani* were found in minimum 1-2 meters away from solfataras (Figure S1) where the concentration of hydrogen sulfide gas was mostly 2-4 ppm (Table S1). We observed fluctuations between zero to ten ppm during the 30 minutes of sampling in Hakkoda.

### Searching for novel symbiotic partners

Whole genomes of the mycobiont, *C. vulcani*, and the photobiont, *Asterochloris* sp., were determined using independent cultures isolated from samples collected in Noboribetsu. The *de novo* assembly of 80.1 M and 144.5 M Illumina Hiseq paired-end reads (150 bp) presented fungal and algal genomes, size and N50 of which are 41.9 Mb and 66.7 Kb, and 55.1 Mb and 149.8 Kb, respectively (Table 1). Completeness of the fungal and algal genomes indicated by Benchmarking Universal Single-Copy Orthologs (BUSCO) assessment was 96.7 % (Ascomycota set) and 86.7 % (Chlorophyta set), respectively.

**Table 1.** Genomic information of the three main symbiotic partners

| Species | Genome size (Mb) | No. of contigs | N50 (Kb) | BUSCO score (%) |
| --- | --- | --- | --- | --- |
| *Cladonia vulcani* | 41.9 | 5523 | 66.7 | 96.7 |
| *Asterochloris* sp. | 55.1 | 990 | 149.8 | 86.7 |
| LFB_Cvu01 | 2.5 | 112 | 26.8 | 52.7 |

In order to identify unknown symbiotic partners of *C. vulcani*, we performed metagenomic sequencing of in total 27 samples from the five localities with at least three biological replicates per locality (Table S2). The metagenomic sequencing yielded 50-100 million 125 bp or 150 bp paired-end reads (Table S2). The metagenomic data were searched for genomic sequences of unknown symbiotic partners. We first performed a draft metagenomic *de novo* assembly using three replicates collected in Noboribetsu. Then we mapped the metagenomic reads to the assembled contigs and plotted the number of mapped reads against the contig length for each sample (Figure 2). The plots of 27 samples clearly showed two distinct lines with steep slopes consisting of high-coverage contigs. BLASTN searches of the high-coverage contigs against the NCBI nr/nt database and the mycobiont genome revealed that the contigs aligned in larger slope among the two are the sequences of the mycobiont, whereas those aligned in smaller slope are highly similar to bacteria of the genus *Achidiphilium*.

### Identification of the third symbiotic partner

We recovered the genome of the *Acidiphilium*-like bacterium that ubiquitously and abundantly exists in all *C. vulcani* metagenomes by assembling consensus sequences of the metagenomic short- and long-reads mapped to the high coverage bacterial contigs and the CDS of *Acidiphilium multivorum* AIU301. The assembly generated 112 contigs, 2.5 Mb in total length and N50 of 26.8 Kb (Table 1). The BUSCO score using the gene set of *Rhodospirillales* was 52.7 %. BLASTN searches indicated that the contigs belong to a novel bacterium [from here on referred to as LFB_Cvu01 (LFB: Lichen Forming Bacterium)] showing high similarity to *Acidiphilium* bacteria.

Phylogenetic trees of the mycobiont and LFB_Cvu01 were constructed using single-nucleotide polymorphism (SNP) sites extracted from the alignment data of the metagenomic short-reads mapped to the genomes (Figure 3). Despite the wide geographical range of the sampling localities (distributed across the three largest islands of Japan, ca. 1,500 km apart in maximum distance), all 27 metagenomes contain reads of LFB_Cvu01 that were mapped to the assembled genome with high similarities (≥ 90 % identify) and coverages (60.63 in average, Table S3). Replicates from the same locality cluster together both in the mycobiont and LFB_Cvu01 trees.

**Figure 3.**
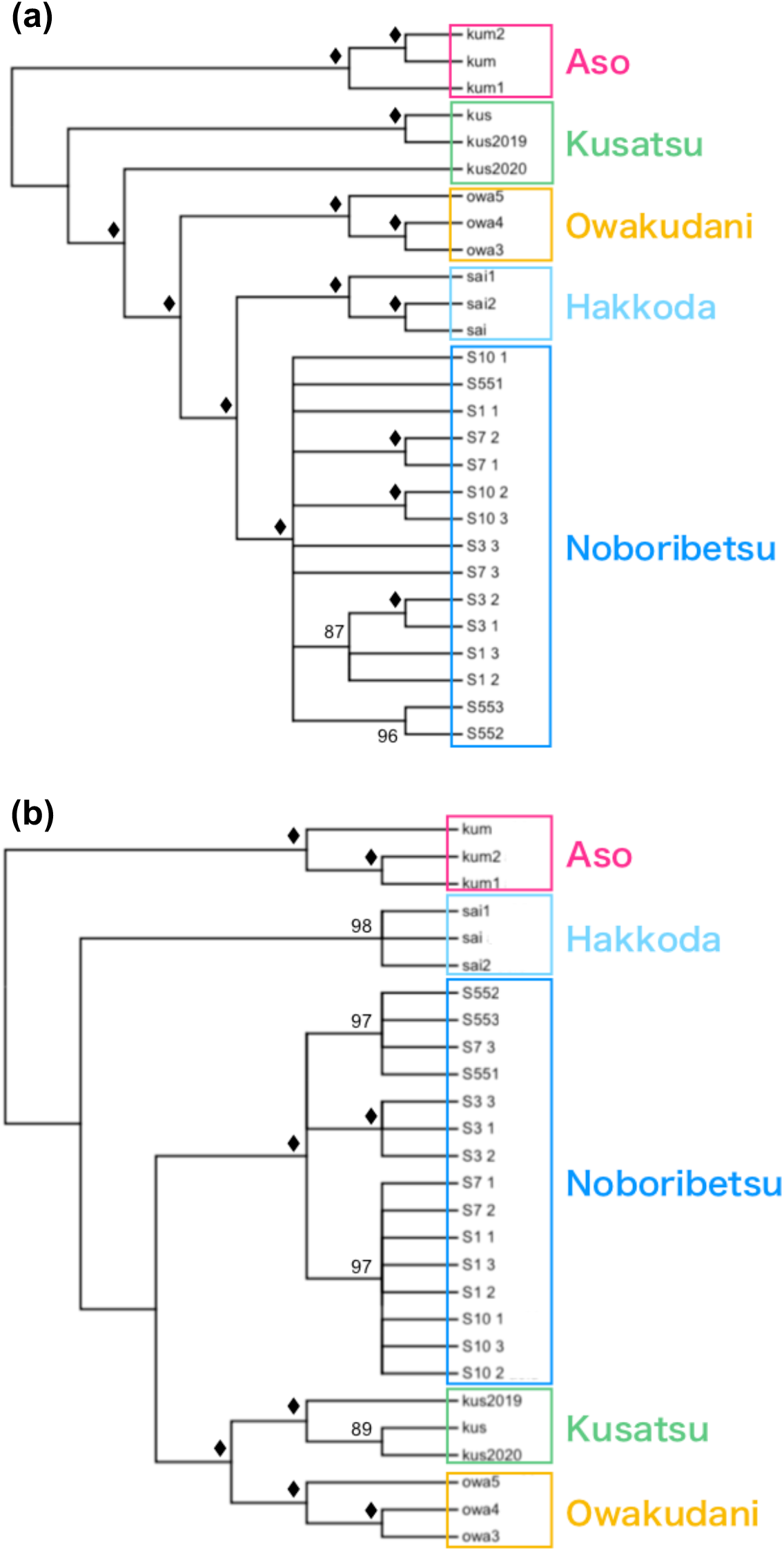
Phylogenetic trees show local clustering of the fungus (a) and the bacterium (b) albeit without clear topological similarities. The diamond symbols indicate branches with bootstrap value of 100.

Abundance of LFB_Cvu01 cells in thalli of *C. vulcani* was predicted from the ratio of bacterial to fungal genome coverage reflecting the ratio of genomic copy number. Among the five localities, average ratios of the replicates varied from 43 to 104 %, lowest in Aso and highest in Hakkoda (Figure 4). Local tBLASTX searches against the contigs of LFB_Cvu01 identified sequences similar to enzymatic genes involved in sulfur metabolism (Table S4 and S5) including sulfite oxidase (e-value: 1.7e-77), and thiosulfate dehydrogenase (quinone) (e-value: 3e-106).

**Figure 4.**
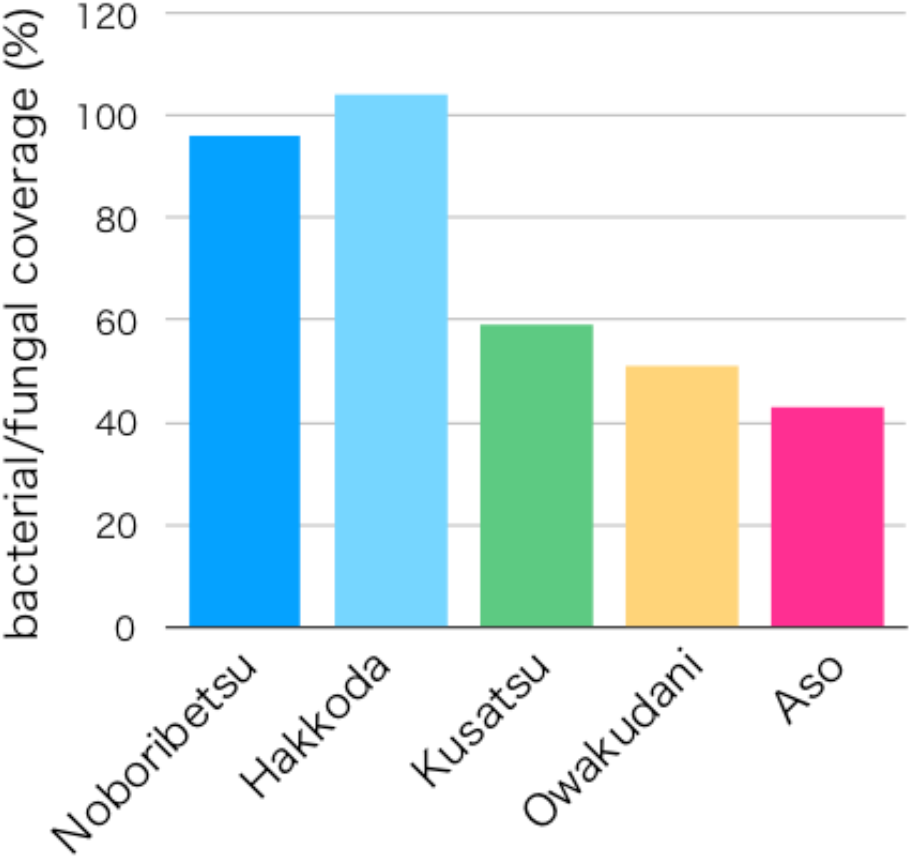
Predicted cellular amounts of LFB_Cvu01 in a thallus of *C. vulcani* relative to the mycobiont in each locality.

### LFB_Cvu01 in ambient soils

Distribution of LFB_Cvu01 in nearby soils was also examined by metagenomic sequencing. The soils were collected 1-2 meters away from three distinct *C. vulcani* populations in Noboribetsu. For each soil sample, 0.00-0.02 % of 37-40 million reads longer than 100 bp were mapped to the genomes of the fungus, alga, and bacterium, respectively.

## Discussion

*Cladonia vulcani* is described as a volcano lichen for its occurrence in volcanic areas especially near solfataras actively emitting toxic sulfurous gases such as sulfur dioxide and hydrogen sulfide (19, 42, 43). Although correlation between sulfurous gases and *C. vulcani* has been indicated (19) the type of gas that affects *C. vulcani* habitation has remained inconclusive. In our study, we detected hydrogen sulfide gas in all sampling locations, whereas sulfur dioxide gas was hardly detected in Kusatsu according to previous reports (44, 45). Therefore it is highly likely that hydrogen sulfide gas is the key factor for the growth of *C. vulcani*.

Hydrogen sulfide gas is highly toxic to most organisms and exposure to high concentrations (more than 1000 ppm) can cause immediate unconsciousness or death in humans. In the sampling locations, *C. vulcani* populations were mostly found in slightly uphill areas 1-2 meters away from solfataras. Although the highest concentration we recorded by the *C*.*vulcani* populations was 10 ppm, devastated surrounding vegetation indicated that the environment is toxic to most of the plants (Figure S2) and *C. vulcani* is likely to have developed mechanisms that enabled adaptation to the extreme environment. Bacteria with abilities that could potentially benefit the lichen symbiosis have been repeatedly isolated from various lichens since the early 20^th^ century (46). The idea of bacteria as an integral part of lichens was corroborated almost a century later by culture-independent approaches that revealed structured composition of lichen-associated bacterial communities (21, 23, 24, 33). While predominance of *Alphaproteobacteria* is a widely shared trend of lichen-associated bacteria at class level comparisons, various biotic and abiotic factors are indicated to be involved in the establishment of lichen-specific communities that are distinct from surrounding environments (20, 21, 23-28, 33). Vertical transmission of bacteria via symbiotic propagules was indicated by previous studies that demonstrated a correlation between the community structure of lichen-associated microbiomes and geographic distance in *Lobaria pulmonaria* (27, 29). Metagenomic and metaproteomic evidence indicated that lichen-associated bacteria are presumed to have functions involved in nutrient supply, stress resistance, detoxification, biosynthesis of secondary metabolites, thallus degradation, and protection against pathogens (25, 30-32). Despite these functions of lichen-associated bacteria that are considered to help lichens to maintain symbiotic association in extreme environments (25, 30, 32), lack of evidence to certify indispensable roles of particular bacterial species in lichen symbiosis has left the significance of lichen-associated bacteria elusive.

Here, we suggest that LFB_Cvu01 is the integral partner of *C. vulcani* which led adaptation to the sulfide-rich extreme environment. LFB_Cvu01 was identified in all 27 *C. vulcani* collected in the five distant localities. In thalli (podetia), LFB_Cvu01 is predicted to exist in 43-104 % of the mycobiont’s cellular amount. This is exceptionally abundant as a single lichen-associated bacterium while the median of individual genus-level bacterial lineages is reported to be 8.3 % in a recent study that examined 437 lichen metagenomes (47). According to the investigation in Noboribetsu, LFB_Cvu01 was not detected in the nearby soils and likely to reside exclusively within *C*.*vulcani* thalli. These tight and specific relations indicate that LFB_Cvu01 is not a mere associate of *C. vulcani* but the essential partner of the symbiosis.

BlastN searches of assembled contigs against the NCBI database indicated that LFB_Cvu01 is similar to bacteria of the genus *Acidiphilium*, some species of which were reported to have sulfur-oxidizing ability (48, 49). Sulfur-oxidizing bacteria can use sulfide or various reduced inorganic sulfur compounds as sources of energy to synthesize organic compounds. Deep-sea hydrothermal vent ecosystems are known to rely on organic compounds fixed by sulfur-oxidizing bacteria (3, 13, 14, 50, 51). Furthermore, sulfide consumption by symbiotic sulfur-oxidizing bacteria contributes to the protection of invertebrate hosts from sulfide-rich vent fluids (10, 11).

The identification of genomic sequences similar to genes involved in sulfur metabolism indicated sulfur-oxidizing ability of LFB_Cvu01. Sulfite oxidase catalyzes a reaction producing sulfate which is the final oxidation product in sulfur oxidation (52). Thiosulfate dehydrogenase (quinone) catalyzes a reaction producing tetrathionate from thiosulfate (53), a common substrate of sulfur-oxidizing bacteria (54-58). Tetrathionate is produced as an intermediate during the oxidation of thiosulfate via “tetrathionate intermediate (S_4_I) pathway” reported from *Beta*-, *Gamma*-, and some *Alphaproteobacteria* (57, 59). Unfortunately we couldn’t identified any complete gene set of known sulfur oxidation pathways in the assembled LFB_Cvu01 contigs. This may be due to the incomplete assembly of the bacterial genome indicated by the low BUSCO score.

Taking the above characteristics of LFB_Cvu01 into account, we hypothesize similar mechanisms as deep-sea vent symbioses in *C. vulcani*. The chemosynthetic activity of the symbiotic bacterium in a thallus provides organic compounds required in the symbiosis, while eliminating sulfide that is toxic to the fungal and algal partners (Figure 5). The establishment of tripartite symbiosis among the fungus, the alga, and the bacterium may have enhanced the fitness of *C. vulcani* as a symbiotic unit in the extreme environment and could have driven its ecological success.

**Figure 5.**
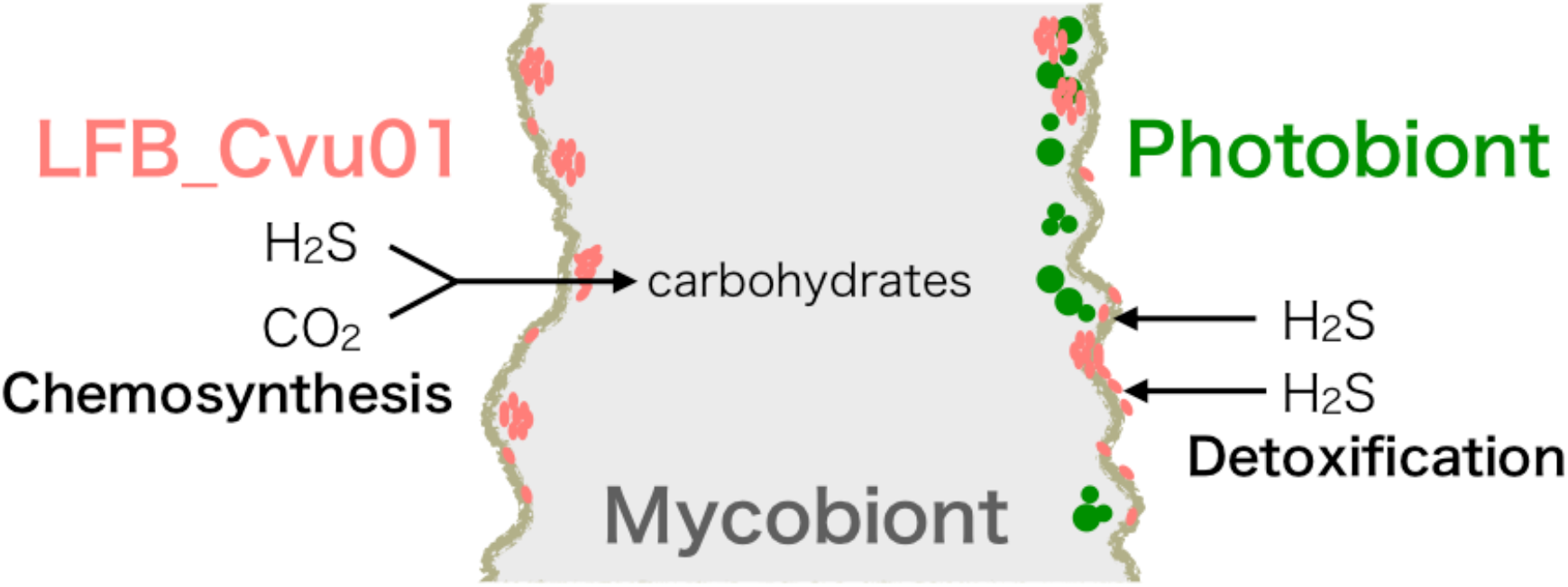
A predicted model of the tripartite symbiosis in *Cladonia vulcani*. The symbiotic bacterium LFB_Cvu01 may contribute in production of carbohydrates and detoxification of hydrogen sulfide by chemosynthesis.

To the best of our knowledge, this could be the first report of chemosynthetic symbiosis in terrestrial geothermal environments that also infers the significant role of a single bacterium species in lichen symbiosis for the first time. Although the complete genome of LFB_Cvu01 is needed to understand its full metabolic capacity, *C. vulcani* will become a model to understand tripartite symbiosis of lichens and terrestrial chemosynthetic symbiosis.

## Acknowledgements

This work was supported by JSPS KAKENHI Grant Numbers JP21H02547 and JP22K15172. We acknowledge Nature Conservation Center Hakone Branch Office and Hakoneonsenkyokyu Co., LTD. for their kind support on our sampling in Owakudani.

