## Supplementary figures for "The third symbiotic partner of the volcano lichen *Cladonia vulcani* Savicz drove adaptation to an extreme environment"

### Supplementary Figure S1

A population of *Cladonia vulcani* (bottom right) covering the ground near a volcanic steam vent (upper left) emitting gas that contains high level of hydrogen sulfide.

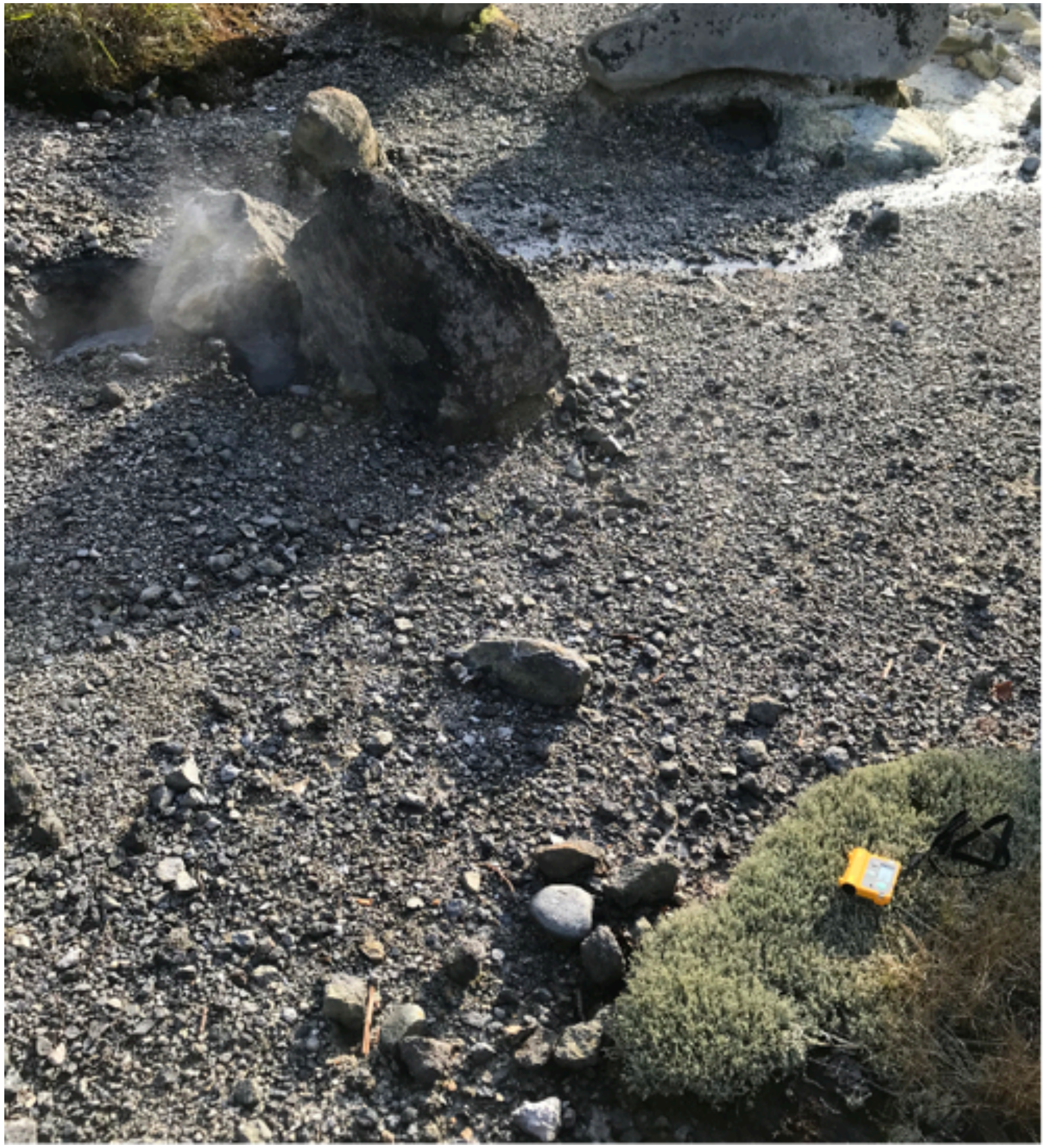

### Supplementary Figure S2

Devastated vegetation around newly formed solfataras. *Cladonia vulcani* is covering the ground cleared of trees and plants (bottom right).

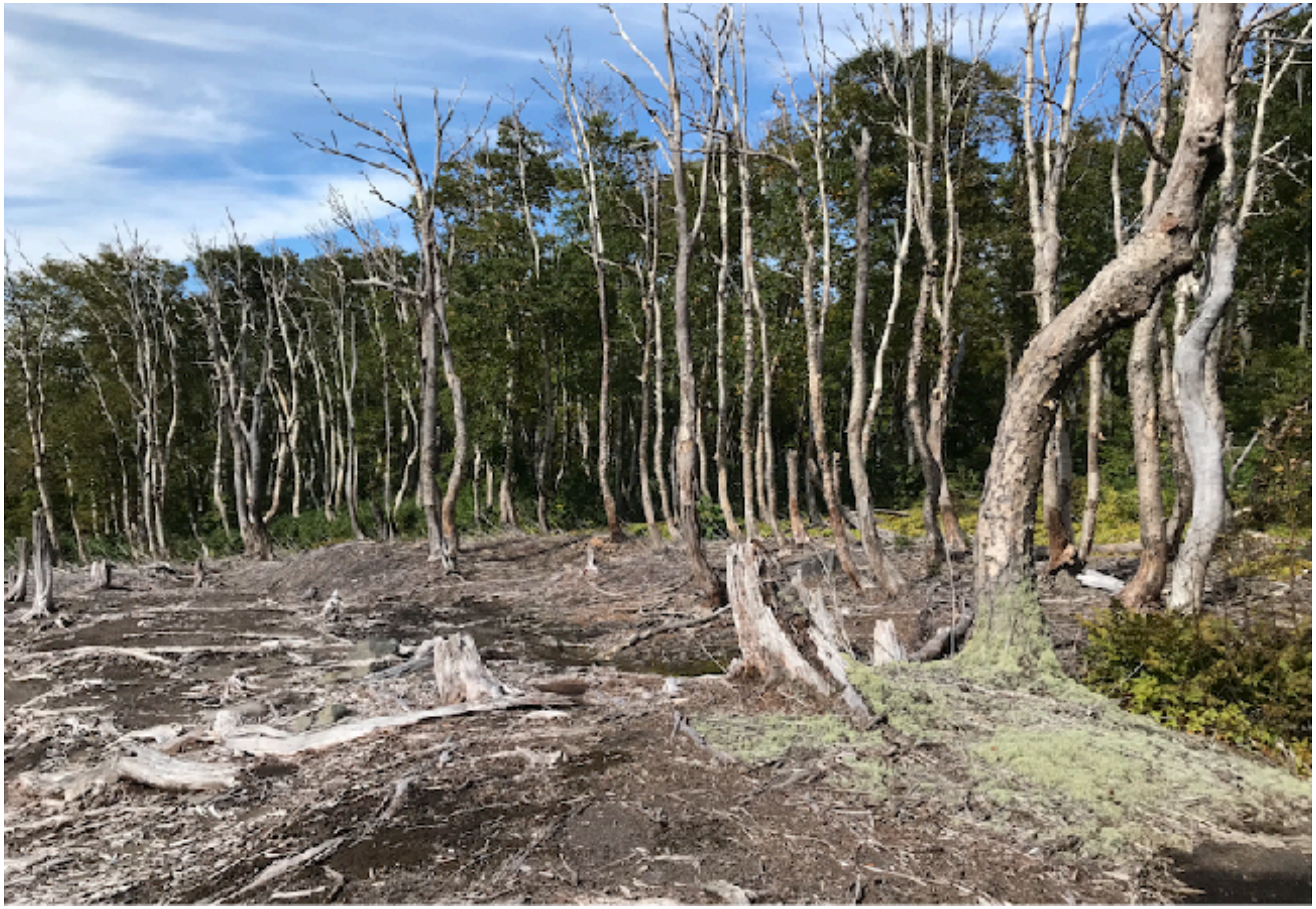
